# Comparative cistromic analysis reveals a shift in tamoxifen-associated endometrial tumor ER*α* binding toward sites less frequently shared by progesterone receptors

**DOI:** 10.1101/2025.09.29.679293

**Authors:** Luciana Ant, Nicolás Bellora, Carina Fariñas, Alejandro La Greca, François Le Dily, Patricia Saragüeta

## Abstract

Tamoxifen, a selective estrogen receptor modulator (SERM) widely used in breast cancer treatment, paradoxically increases endometrial cancer risk. Although progesterone receptor (PR) shares estrogen receptor alpha (ERa) binding sites in Ishikawa endometrial adenocarcinoma cells, that could explain in part progestin regulation of estrogen effects, whether long-term tamoxifen exposure alters this ERa–PR interplay in endometrial tumors remains unknown.

Given the absence of endometrial tumor PR cistrome data, we compared ERa cistromes from tumors of users and non-users with Ishikawa cells ERa and PR cistromes to investigate how long-term SERM exposure might affect ERa binding signatures. Here, we show that non-user ERa binding exhibits greater similarity to the PR cistrome than tamoxifen users. These shared ERa-PR binding sites are not enriched in estrogen response elements (EREs) but in AP-1 and SOX motifs, suggesting indirect or tethered ERa recruitment, and are associated with genes involved in inflammatory pathways, such as MYC. In contrast, tamoxifen exposure appears to shift ERa binding toward PR-independent regions enriched in ERE motifs, and that could regulate early estrogen response pathways genes, including the oncogene NRIP1. Our findings suggest that exposure to tamoxifen may redirect ERa binding toward genomic regions responsive to estrogen, whereas ERa binding in non-users retains features associated with PR-occupied regulatory sites.

These results provide bases for further investigation about the role that SERM treatment could play in remodeling the endometrial tumor epigenetic landscape and point to the hypothesis that this remodeling may involve altered ERa–PR regulatory interplay, potentially contributing to endometrial tissue dysregulation.

**Significance:** This study suggests that long-term tamoxifen exposure may alter ER*α* regulatory patterns in endometrial tumors, shifting binding away from PR-associated regions toward estrogen-responsive genomic sites. These differences raise the possibility that tamoxifen reshapes the endometrial tumor epigenetic landscape by modifying ER*α*–PR interplay, with potential consequences for gene regulation and tissue homeostasis.

## Introduction

Endometrial cancer (EC) is the sixth most common malignancy worldwide (1). Epidemiological trends indicate a rapid rise in EC incidence (1, 2), with mortality rates increasing by 1.8% annually since 2010 (1, 2). Historically, EC has been classified in two categories: Type I or Type II (3). Type I EC, which are more prevalent and generally associated with a favorable prognosis, are linked to unopposed estrogen exposure and typically consist of grade I or II endometrioid adenocarcinomas. In contrast, Type II ECs are considered hormone-independent and tend to be more aggressive, encompassing grade III endometrioid adenocarcinomas, serous, clear cell, undifferentiated carcinomas, and carcinosarcomas (4, 5).

In hormone-dependent Type I EC, the interplay between estrogen and progesterone signaling-mediated by estrogen receptor alpha (ERa) and progesterone receptor (PR), respectively-plays a crucial role in tumor development and progression. Estrogen promotes the proliferation of the endometrial glandular epithelium-the primary cell type involved in EC-whereas progesterone antagonizes these estrogenic effects (6). An imbalance between estrogen and progesterone exposure has been associated with endometrial carcinogenesis (7). Specifically, prolonged exposure to unopposed estrogen can result in endometrial hyperplasia and overgrowth, a recognized precursor to malignancy (8, 9). Progesterone exerts a protective effect by inhibiting estrogen-mediated proliferation and promoting cellular differentiation (6). Notably, the loss of PR expression has been correlated with increased proliferation in both ER-positive and ER-negative endometrial tumors (10).

Estrogens, particularly 17*β*-estradiol (E2), are essential for the growth and differentiation of various tissues, including the endometrium, where they promote proliferation through ERa and ER*β*. ERa modulates gene expression via several mechanisms, the most classical of which involves direct binding to estrogen response elements (EREs) within chromatin, regulating target genes such as the progesterone receptor gene (PGR) (11). In addition, ERs can influence transcription through ERE-independent pathways, wherein DNA binding is facilitated by interactions with other transcription factors, such as Fos and Jun (11). Beyond these genomic mechanisms, estrogens can also act through non-genomic pathways, such as signaling via the G protein coupled estrogen receptor (GPER/GPR30) (12). In this study, we focus specifically on genomic ERa- and PR-mediated chromatin binding.

Progesterone mediates its biological effects through two main isoforms of the progesterone receptors, PRA and PRB, which are transcribed from distinct promoters of the PGR gene (13). Upon ligand binding, progesterone receptors modulate gene expression by interacting with progesterone response elements (PREs) in the genome, thereby regulating the transcription of downstream target genes.

Selective estrogen receptor modulators (SERMs), such as tamoxifen, are commonly used in treatment of estrogen receptor-positive breast cancer. Tamoxifen exerts its effects by binding to the ligand-binding domain of ERa. While it functions predominantly as an ERa antagonist in breast tissue, inhibiting estrogen-driven proliferation (14, 15), its activity is highly tissue-specific. In the endometrium, tamoxifen paradoxically acts as an ERa agonist, promoting epithelial proliferation and increasing the risk of EC (16–18). In addition, tamoxifen has also been shown to act as an estrogen agonist through non-genomic pathways such as GPER (19), through which can induce cell migration in EC (20). At the genomic level, studies using the Ishikawa EC cell line have demonstrated that tamoxifen modulates gene expression through ERa-dependent mechanisms (21). Genome-wide analyses further revealed that the ERa cistromes of tamoxifen-associated endometrial tumors overlap significantly with those of breast cancer (22). Moreover, tamoxifen-associated endometrial tumors exhibit a distinct ERa DNA-binding profile compared to spontaneous cases, with these altered binding patterns correlating with differential gene expression (23).

In breast cancer, PR modulates ERa activity by influencing its chromatin binding dynamics and downstream gene expression (24). Studies in T47D mammary cancer cells have identified hormone control regions (HCRs), which harbor both ERa and PR binding sites and physically interact with promoters of hormone-regulated genes within hormone-responsive topologically associating domains (TADs) (25, 26). In contrast, the interplay between ERa and PR in EC remains poorly characterized. Recent work by La Greca et al. (27) demonstrated that estrogen priming induces the formation of progestin-dependent PR binding sites in Ishikawa EC cells. These sites are frequently located within ‘progestin control regions’ (PgCRs), a subset of HCRs situated in TADs, involving coordinated binding of ERa, PR, and the transcription factor PAX2 to regulate gene expression. Notably, PgCRs are characterized by an open chromatin conformation and are associated with the regulation of genes implicated in endometrial carcinogenesis.

While it is established that prolonged tamoxifen treatment is associated with a distinct ERa binding profile in endometrial tumors compared to those not associated with tamoxifen, the relationship between these altered ERa binding sites and the regulatory landscape of the PR remains uncharacterized. The interplay between ERa and PR is critical in hormone-dependent EC, with PR having the capacity to modulate ERa’s effects (27). Previous work has high-lighted that ERa’s positioning is a key determinant in the development of hormone-dependent EC (23). Therefore, this study aims to investigate tamoxifen-induced changes in the ERa cistrome related to PR binding sites in EC cells. We hypothesize that tamoxifen disrupts the co-regulatory relationship between ERa and PR, leading to a distinct oncogenic program. To test this, we conducted a comparative genomic analysis integrating publicly available ERa ChIP-seq data from endometrial tumors of tamoxifen users and non-users with our previously published PR and ERa cistromes from the Ishikawa EC cell line. This approach allowed us to characterize differences in ERa binding patterns between tamoxifen-associated and non-associated ECs in the context of PR-associated regulatory regions, providing evidence for a potential role of altered ERa–PR regulatory interplay in tamoxifen-associated endometrial cancer.

## Methods and Materials

### Patient Material

Public available patient material was used for microarray and ChIP-seq analysis. The data, published by Droog *et al*. (23), was derived from the TAMARISK study and consists of endometrial samples of anonymized patients, who signed an informed consent. The TAMARISK study was performed in accordance with the Code of Conduct of the Federation of Medical Scientific Societies in The Netherlands and has been approved by the local medical ethics committee of The Netherlands Cancer Institute. According to the original study, tamoxifen users and non-users were comparable with respect to the available clinicopathological characteristics, including age, tumor grade, FIGO stage, and MSI status (23). Detailed information on sample collection and patient characteristics can be found in the original publication (23).

### ChIP-seq Data Processing

Fastq sequence reads from GEO database (see Data Availability) high-quality sequencing reads were aligned to the human reference genome (hg38, UCSC) using STAR v2.7.11 (28), with alignment parameters set to a maximum intron size of 1 and end-to-end alignment enabled. For initial quality control and sample clustering, reads for each sample were aligned independently. Subsequently, samples were stratified based on tamoxifen exposure (tamoxifen users vs. non-users) and reads within each group were merged into a single BAM file for downstream analysis (29, 30). Peak calling was performed using MACS2 v2.2.6 (31) with default parameters, employing the corresponding input controls to identify regions of significant signal enrichment. For all the datasets, only binding sites with an FDR *≤* 0.05 were included. Venn diagrams illustrating peak sets overlaps were generated using the venn function from the Intervene package (32). Principal Component Analysis (PCA) and heatmap visualizations were generated using deepTools (33). Specifically, computeMatrix was used to calculate signal intensities, plotHeatmap for heatmap generation, multiBigwigSummary for region-wise signal quantification, and plotPCA for dimensionality reduction. Genomic overlap and distance between ERa and PR binding sites in Ishikawa cells were assessed using the ‘intersectBed’ and ‘closest’ utilities from bedTools (34).

### RNA-seq Data Processing

Differential gene expression analysis was conducted using the DESeq2 package v1.44.0. (35). Genes exhibiting an absolute log_2_ fold change (|log2FC|) *≥* 1 and a Benjamini-Hochberg adjusted p-value (padj) *≤* 0.05 were considered differentially expressed. Expression levels, quantified as RPKM (Reads Per Kilobase of transcript, per Million mapped reads) and TPM (Transcripts Per Million) were visualized using custom Python scripts employing the seaborn library (36).

### Gene association with Binding Sites and Gene Ontology Enrichment analysis

Binding sites were associated with genes using two complementary approaches. The first one was done using GREAT algorithm (37, 38). This algorithm first assigns every gene a regulatory domain and then associates each genomic region with all genes whose regulatory domain it overlaps. The association was done using the default “basal plus extension” setting. With this setting, each gene is assigned a basal regulatory domain extending 5 kb upstream and 1 kb downstream of the transcription start site (TSS), independent of the regulatory domains of neighboring genes. The distal regulatory domain extends bidirectionally from the basal domain until reaching the basal domain of the nearest gene, with a maximum extension of 1 Mb in either direction. Binding sites overlapping either the basal or distal regulatory domain of a gene were therefore associated with that gene. The second approach to find genes associated with binding sites was performed by selecting the TADs obtained from Ishikawa Hi-C (see Hi-C Data Processing section) that comprised binding sites of interest and retrieving all the genes contained in that TAD. Gene Ontology (GO) enrichment analysis was performed using the GSEApy package (39), a Python-based implementation of GSEA and a wrapper for the Enrichr database. The enrichr function was employed to query the following libraries: ‘BioCarta_2016’, ‘KEGG_2021_Human’, ‘Reactome_2022’, ‘MSigDB_Hallmark_2020’, and ‘Else-vier_Pathway_Collection’. Any pathway enrichment with an associated padj value *≤* 0.01 was considered significantly enriched.

### Microarray Data processing and Analysis

Microarray gene expression data from endometrial cancer samples derived from 47 tamoxifen users and 64 non-users was retrieved from GEO (see Data Availability). The microarray data was provided as log_2_-transformed expression ratios derived from dye-swap experiments, as described in the original study (23). Differential gene expression between tamoxifen users and non-users was assessed using the Limma R package (40). No gene was found significantly differentially expressed (padj *≤* 0.05) between tamoxifen users and non-users. Gene set enrichment analysis (GSEA) was subsequently performed with GSEApy package (39) gsea function, using the expression data without applying a differential expression significance cutoff or gene pre-filtering, with the following parameters: permutation type set to “phenotype”, 1000 permutations, and “difference of class means” as the scoring method. Enriched terms with an adjusted padj *≤* 0.1 were considered significant.

### Hi-C Data Processing

Previously filtered Hi-C datasets from (27), excluding lowquality reads, were further processed using the Juicer pipeline (41) with the human reference genome hg38. After individual sample contact maps were generated, contact matrices from biological replicates were merged by summing the contact counts between corresponding genomic loci, resulting in a combined contact map. Topological Associated Domains (TADs) were identified using the Arrowhead algorithm, and chromatin loops (i.e., locally enriched contacts) were detected using the HiCCUPS algorithm, both implemented within the Juicer framework with default settings. For visualizations, Hi-C contact matrices were normalized using Square Root Vanilla Coverage.

### Virtual 4C

Virtual 4C profiles were generated using the capture Hi-C (chic) module of HiCExplorer (42). Reference viewpoints were defined as a 1-kb window centred on the transcription start site (TSS *±* 500 bp) of the NRIP1 and MYC promoters, based on GRCh38 gene coordinates retrieved from GeneCards. Hi-C replicate matrices with a 10-kb resolution were merged and normalized using Knight-Ruiz (KR) and used as input for each condition (T0, E2 and R). HiC-Explorer function ChicViewpoint was used to compute the interaction profile of each viewpoint and condition within a window of 550 kb upstream and downstream of the reference point. Relative interaction values were then extracted from chicViewpoint output, converted to bedGraph format, and visualized alongside the corresponding Hi-C matrices using pyGenomeTracks (42, 43).

### Motif enrichment analysis

FASTA sequences corresponding to ERa binding sites from tamoxifen non-users overlapping Ishikawa PR binding sites, Ishikawa ERa binding sites not overlapping PR sites, and Ishikawa ERa binding sites overlapping PR sites were analyzed separately using Simple Enrichment Analysis (SEA; MEME Suite v5.5.9) (44). SEA was performed against the human Catalog of Inferred Sequence Binding Preferences (CIS-BP) 2.00 motif database (1065 motifs) using a second-order Markov background model and center alignment. A permissive enrichment threshold of 10000 was used to retain all motifs reported by SEA for subsequent filtering. To reduce redundancy among enriched motifs, motifs were clustered according to the genomic overlap of their predicted motif matches (Figure 1—Figure supplement 3), following the approach described by Bellora et al. (45). Motifs that reached a False Discovery Rate (FDR) *≤* 0.05 in at least one of the three groups were included in the clustering analysis (204 motifs). ZNF motifs were excluded from the primary analysis because of their large number and limited functional annotation, leaving 139 motifs. An additional clustering done including ZNF motifs produced similar clustering patterns. Similarity was calculated as in Bellora *et al*. (45). Briefly, two motif matches were considered overlapping when they shared at least 3 bp, irrespective of strand. For each pair of motifs A and B, similarity was calculated from the proportion of sites of each motif overlapping at least one site of the other motif:

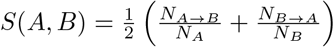

where *N*_*A*_ and *N*_*B*_ are the total numbers of predicted sites for motifs A and B, respectively, and *N*_*A→B*_ and *N*_*B→A*_ are the numbers of sites of motifs A and B, respectively, that overlap at least one site of the other motif. The resulting similarity values were converted to distances as *D*(*A, B*) = 1 *− S*(*A, B*). Hierarchical clustering was implemented with the scipy.cluster.hierarchy module (SciPy version 1.12.0), using average linkage, and a distance cutoff of 0.6 was used to define motif clusters. Of the 139 motifs included in the analysis, 37 did not cluster with another motif at this cutoff and therefore remained as individual motifs.

**Figure 1.**
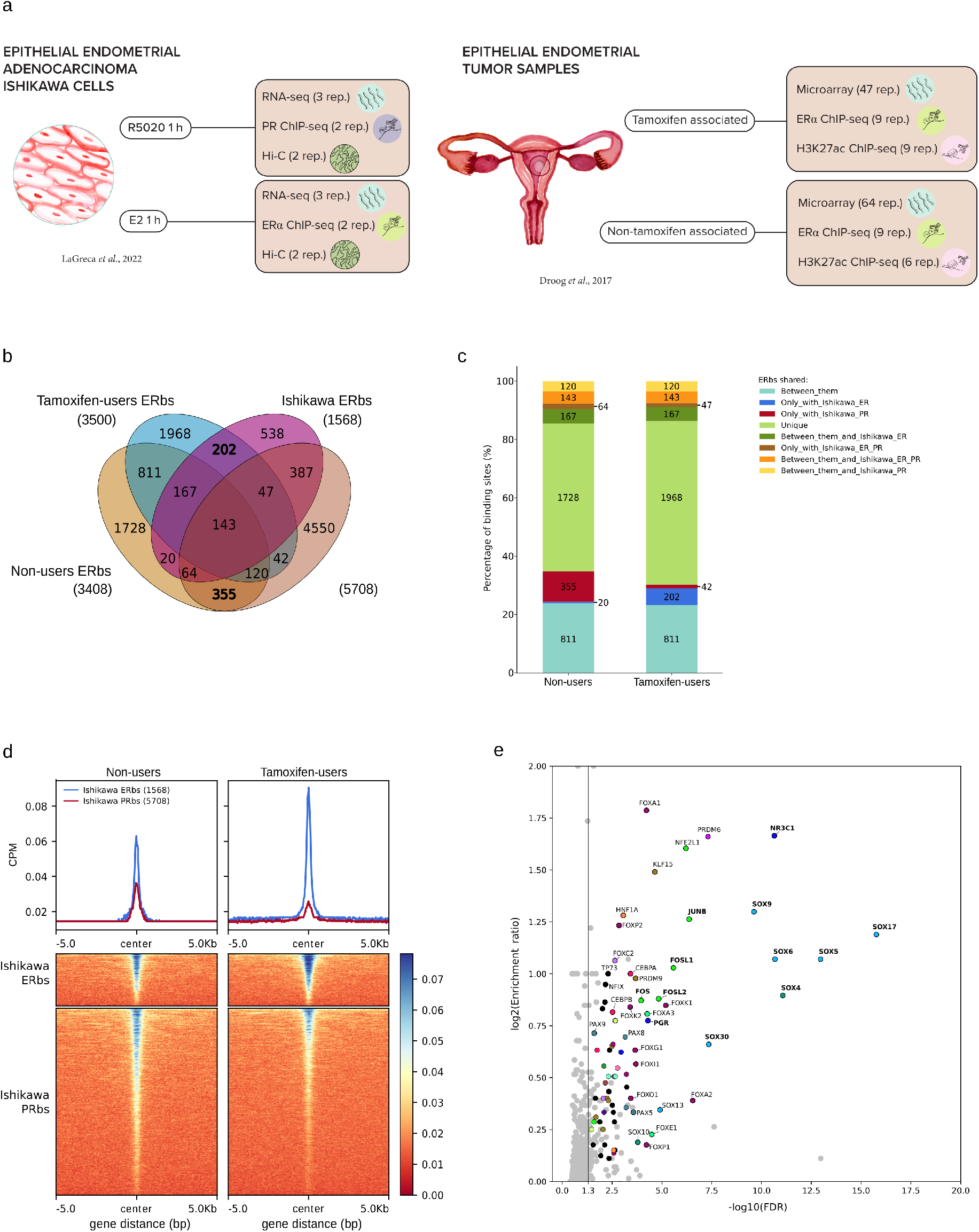
Comparative analysis of ERa and PR binding profiles in endometrial tumors and Ishikawa cells. **a.** Schematic overview of the experimental set up involving endometrial tumors from tamoxifen users and non-users, as well as Ishikawa endometrial adenocarcinoma cells. **b**. Venn diagram showing the overlap between PR binding sites (PRbs) and ERa binding sites (ERbs) in Ishikawa cells, along with ERbs identified in tamoxifen users and non-users. For all datasets, only binding sites with an FDR *≤* 0.05 were included. The total number of binding sites (bs) for each group is indicated in parentheses. **c**. Percentage of ERbs in tamoxifen users and non-users, categorized by their sharing pattern: shared between users and non-users (Between_them); shared only with Ishikawa ERa (Only_with_Ishikawa_ER) or Ishikawa PR (Only_with_Ishikawa_PR); unique to each group (Unique); shared between users, non-users, and Ishikawa ERa (Between_them_and_Ishikawa_ER); shared only with both Ishikawa ERa and PR (Only_with_Ishikawa_ER_PR); shared across all groups (Between_them_and_Ishikawa_ER_PR); and shared between users, non-users, and Ishikawa PR (Between_them_and_Ishikawa_PR). **d**. ERa ChIP-seq signal (CPM) in tumors from tamoxifen users and non-users. Signal was quantified within a 10 kb window centered on Ishikawa ERbs and PRbs (*±*5 kb). Top panel: Median CPM signal across all Ishikawa ERa and PRbs. Bottom panels: Signal intensity for each individual region. **e**. Scatter plot of the log2 enrichment ratio of each motif against its statistical significance (log10(FDR)). Enrichment ratios were calculated with Simple Enrichment Analysis (see Materials and Methods). The vertical line marks the FDR = 0.05 threshold. Motifs with a significant enrichment (FDR *≤* 0.05) are colored, with motifs belonging to the same cluster sharing the same color (Figure 1—Figure supplement 3) and motifs that did not cluster with any other motif shown in black. Motifs that were not significant are shown in light gray. ZNF motifs were excluded. Labeled motifs were selected manually for visualization, without a predefined selection criterion. Bold labels indicate mot1if3s discussed in the text.

### Endometrial Cancer Samples (TCGA)

RNA-seq gene expression data (STAR-Counts workflow, TSV format) for human endometrial cancer samples (n=394) were obtained from The Cancer Genome Atlas (TCGA) Uterine Corpus Endometrial Carcinoma (TCGA-UCEC) open-access project. The cohort was filtered to include primary tumor tissue from the endometrium (Primary

Diagnosis: endometrioid adenocarcinoma, NOS; Disease Type: adenomas and adenocarcinomas) with no reported synchronous malignancy. Each sample was annotated with its corresponding FIGO stage. Differential expression analysis of NRIP1 across FIGO stages was performed using pairwise comparisons with Tukey’s Honestly Significant Difference (HSD) post-hoc test, implemented via the statsmodels Python package (46). An adjusted p-value (padj) *≤* 0.05 was considered statistically significant. For correlation of gene expression analysis, TPM counts were transformed to log2(TPM), to reach a normal distribution. A Pearson partial correlation was calculated, with age of the subject and FIGO stage of the sample as covariates. Spearman correlation yielded a similar correlation value. For overall survival (OS) analysis, following a protocol similar to Li *et al*. (47), mRNAs with *≥* 1 FPKM in at least 10% of the samples were retained (n=17180). Pathway enrichment scores were calculated via ssGSEA using the GSVA (48) and msigdbr (49) R packages, and standardized to z-scores across samples. Using the overall cohort median z-score as a cutoff, patients were divided into high (z-score *≥* median) and low (z-score < median) pathway expression groups. Subsequently, they were analyzed across: all FIGO stages, early (Stage I–II), and advanced (Stage III–IV) disease. Kaplan-Meier curves and log-rank tests were generated using the survival (50) and survminer (51) R packages.

## Results

### A comparative framework to relate ERa binding in tamoxifen-associated endometrial tumors to Ishikawa ERa and PR cistromes

To investigate how tamoxifen alters the chromatin landscape of ERa in relation to PR signaling, we conducted a comparative analysis of ERa binding sites from endometrial tumors of tamoxifen users and non-users. Due to the absence of publicly available PR binding data from endometrial tumors, we employed our previously characterized ERa and PR cistromes from the Ishikawa EC cell line as a reference framework for interpreting the changes in ERa cistrome of users and non-users.

The ERa landscape of tamoxifen user and non-user endometrial tumors (22, 23) were compared to our previously published ERa and PR cistromes generated with an antibody targeting ERalpha and an antibody that detects both PR isoforms respectively, from Ishikawa EC cell line treated with 17*β*-estradiol (E2) or the synthetic progestin R5020 (27) (Figure 1a). In our previous work, we observed that estrogen pre-treatment leads to an increase in the total PR binding sites (PRbs), while retaining 97% of PRbs identified without E2 pre-treatment (Figure 1—Figure supplement 1a). Accordingly, we focused on the PR cistrome in estrogen-primed Ishikawa cells to characterize PR binding within the identified estrogen-responsive context (Figure 1—Figure supplement 1a).

For this comparison, ERa ChIP-seqs raw available data (23) from 6 non-users tumor and 9 tamoxifen users tumors were reprocessed under the same conditions as the ChIP-seq of Ishikawa cell line (Figure 1—Figure supplement 1b). Although the samples were heterogeneous, as expected for clinical derived data, principal component 2 values derived from a Principal Component Analysis (PCA) and hierarchical clustering were able to distinguish the two groups (Figure 1—Figure supplement 1c, d).

To characterize the ERa binding site population in the endometrial tumor samples, we classified sites into three categories: unique to tamoxifen users, unique to non-users, and common to both (Figure 1—Figure supplement 2a). In tamoxifen users, the ERa ChIP-seq signal showed a loss of 2167 binding sites compared to non-users, while tamoxifen treatment contributed to the acquisition of additional 2259 binding sites in tumors (Figure 1—Figure supplement 2a). To associate these sites with enhancer activity, H3K27ac levels were analyzed, revealing that tamoxifen-treated tumors exhibited increased activity at the binding sites gained after treatment (Figure 1—Figure supplement 2b).

### Tamoxifen-associated shifts in ERa binding sites reflect estrogen-responsive, PR-independent regulation

To understand the functional ERa binding landscape in tumor samples, we quantified the overlap with the ERa binding sites (ERbs) and PR binding sites (PRbs) identified in Ishikawa cells. This revealed a distinct pattern between non-users and tamoxifen users (Figure 1b). Ishikawa ERbs shared 3 times more binding sites with ERbs from tamoxifen users than with those from non-users (Figure 1b). Conversely, Ishikawa PRbs exhibited 4.7-fold greater overlap with ERbs from non-users compared to tamoxifen users (Figure 1b). This asymmetry was also evident considering in the percentage of binding sites uniquely shared: 5.8% of all ERbs in tamoxifen users were uniquely shared with Ishikawa ERbs (202 ERbs), while only 1.2% were uniquely shared with Ishikawa PRbs (20 PRbs). In contrast, non-users shared 0.59% of their ERbs exclusively with Ishikawa ERbs (42 ERbs) and 10.4% with Ishikawa PRbs (355 PRbs) (Figure 1c). Furthermore, the ChIP-seq signal intensity for ERa at the binding sites shared with Ishikawa ERa was higher in tamoxifen users than in non-users, whereas non-users showed a stronger ERa signal at sites overlapping with Ishikawa PRbs (Figure 1d).

These results suggest that tamoxifen exposure in endometrial tumors is associated with an ERa chromatin binding profile that more closely resembles the ERa binding land-scape observed in estrogen-treated Ishikawa cells. In contrast, the ERa binding profile of non-tamoxifen users exhibits greater similarity to the PR binding pattern in the same cell line.

To identify transcription factors that could be associated with the 355 ERbs from non-users that overlap Ishikawa PRbs, we performed a motif enrichment analysis on these sequences. Since the motif database contains many motifs with similar sequence specificity, motifs significantly enriched (FDR *≤* 0.05) were clustered according to the genomic overlap of their predicted sites (Figure 1—Figure supplement 3, Materials and Methods). Within these sites, we found a significant enrichment of 76 motifs (FDR *≤* 0.05) (Figure 1e). The motifs for the nuclear receptors NR3C1 (GR) and PGR showed an enrichment ratio of 3.5 and 1.6 respectively. Given the similarity between the response elements of these steroid receptors (Figure 1—Figure supplement 3), the enrichment of the NR3C1 and AR motifs may result from the presence of PGR motifs. Several SOX motifs were also enriched, including SOX4, SOX5, SOX6, SOX9, SOX17 and SOX30. This is of particular interest given the reported SOX motif enrichment in ERa binding specific to endometrial cancer (52). The ESR1 and ESR2 motifs were not enriched in these ERbs. Instead, we found a significant enrichment of the AP-1 cluster (FOS, FOSL1, FOSL2 and JUNB, together with NFE2L1). Since ERa can bind to the chromatin through tethering to Fos/Jun (53), the enrichment of these motifs indicate that ERa could be recruited to these sites through AP-1 rather than by direct binding to an ERE.

### ERa/PR binding in non-user links to inflammatory gene expression

To understand the potential regulatory roles of the 355 unique non-user ERbs that overlap with Ishikawa PR binding, we employed two complementary approaches to associate these regions with nearby genes.

#### GREAT based approach

First, we used the GREAT algorithm (37) using the default “basal plus extension” association rule to assign genes to the ERa binding sites based on their regulatory domains (see Materials and Methods). The genes whose regulatory domains contained one of the 355 unique non-user ERbs overlapping Ishikawa PRbs were selected. Gene ontology (GO) analysis of the associated gene set revealed significant enrichment in the Mucin type O-glycan biosynthesis pathway, Basal transcription factors, and two NOTCH-related pathways: NOTCH receptor signaling and NOTCH signaling deregulation in cancer (Figure 2a and supplementary material). Given the established involvement of the NOTCH pathway in various cancers-including breast, ovarian, and cervical (54)-we assessed potential differential expression of these pathways in the publicly available microarray dataset of endometrial tumors from tamoxifen users and non-users (23). Limma analysis of this dataset for individual differentially expressed genes failed to reveal any significant differences (padj *≤* 0.05)(Figure 2—Figure supplement 1a). Gene set enrichment analysis (GSEA) of “NOTCH signaling deregulation in cancer” did not reveal statistically significant enrichment (FDR > 0.1) (Figure 2—Figure supplement 1b). This could be due to the high variability among the microarray samples and the small size of the gene set (29 genes) compared to the microarray entire gene set (~ 14000 genes). On the other hand, gene expression of NOTCH-related genes in Ishikawa cells treated with ethanol (vehicle), estradiol (E2), or tamoxifen revealed differential expression of several (CCND1, CDKN1A, FURIN, HES1, JAG1, JAG2, MYC, NCOR2, NCSTN, NOTCH1, NOTCH3 and TWIST1) genes within both the “NOTCH receptor signaling” and “NOTCH signaling deregulation in cancer” pathways (Figure 2—Figure supplement 1c) (55).

**Figure 2.**
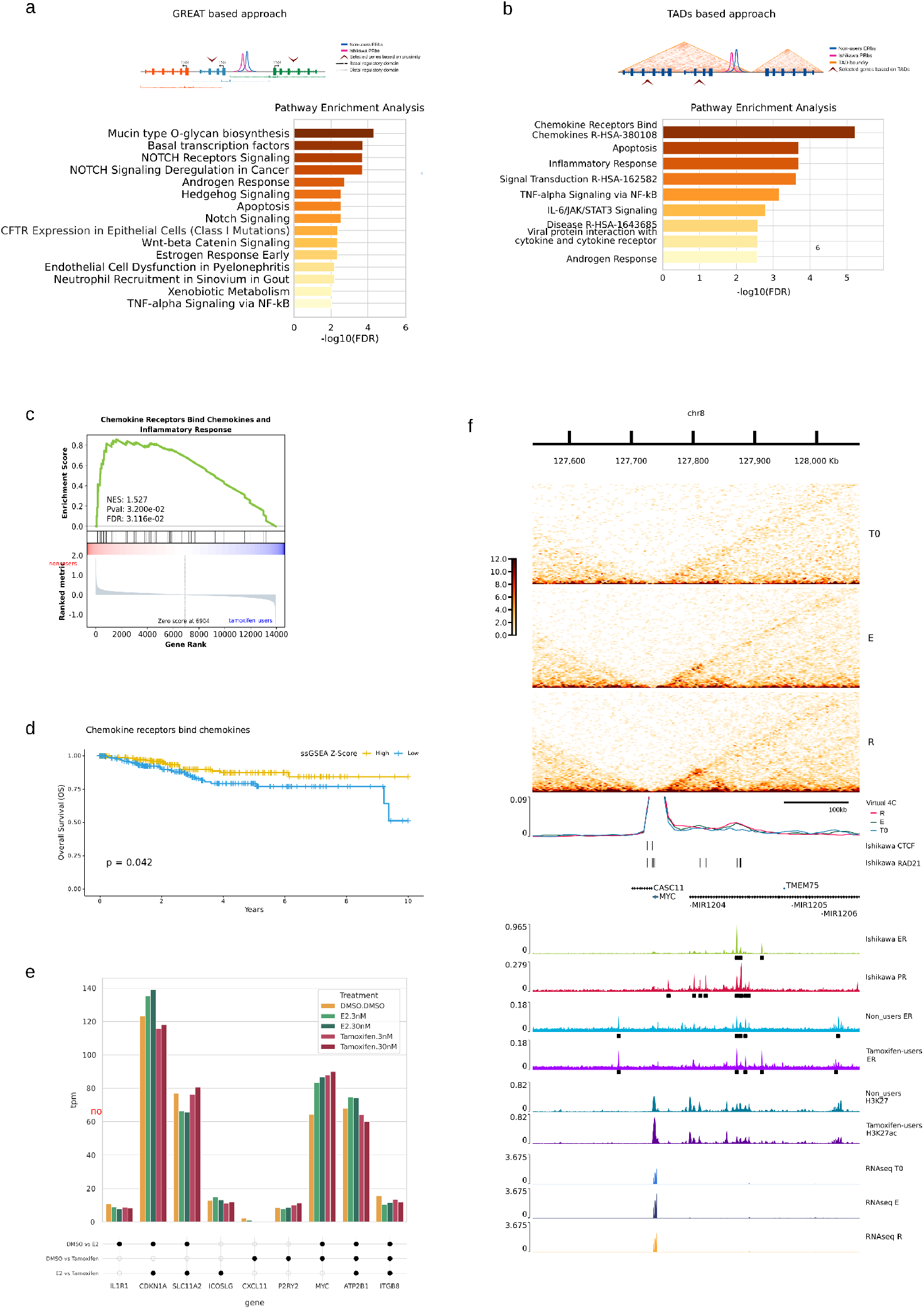
Pathway enrichment analysis of genes associated with PR and non-user ERa co-bound regions. **a.** Top enriched pathways among genes located near Ishikawa PRbs that overlap with ERbs from non-user (*±*5 kb from the transcription start site, TSS) (left). Representation of the GREAT-based approach used to associate binding sites to nearby genes (right). The analysis was performed using GREAT’s default “Basal plus extension” association rule. Under this setting, the basal regulatory domain of each gene extends 5 kb upstream and 1 kb downstream of the TSS, while the distal regulatory domain extends in both directions from the basal domain until it reaches the basal domain of the nearest gene, with a maximum extension of 1 Mb in either direction. The genes whose regulatory domains contained one of the 355 unique non-user ERbs overlapping Ishikawa PRbs were selected. **b**. Top enriched pathways in genes located within topologically associating domains (TADs) that contain at least one Ishikawa PRbs overlapping with an ERbs from non-users (left). Representation of the TADs-based approach used to associate binding sites to nearby genes (right). **c**. Gene set enrichment analysis (GSEA) of gene expression profiles from tamoxifen users and non-users. The analyzed gene set includes Chemokine Receptors Bind Chemokines and Inflammatory Response genes located in TADs containing at least one Ishikawa PRbs overlapping with an ERbs from non-users. The enrichment profile (green) indicates the correlation of gene expression with the ranked expression dataset. Normalized enrichment score (NES) and nominal p-value (nom.p) are reported. **d**. Kaplan-Meier overall survival (OS) curves of endometrial cancer patients (TCGA data) stratified into high (yellow) and low (blue) pathway expression groups, using the median ssGSEA z-score as a cutoff for the Chemokine Receptors Bind Chemokines pathway. Log-rank test p-value is reported. **e**. Expression levels (TPM) of genes within the Chemokine Receptor Bind Chemokines and Inflammatory Response pathways in Ishikawa cells treated with control (DMSO), tamoxifen (3 nM and 30 nM), or estradiol (3 nM and 30 nM) (E2.3 nM and E2.30 nM) (55). Black circles indicate significant differential expression between the corresponding treatment groups (padj *≤* 0.05). **f**. Genomic architecture, epigenetic landscape, and gene expression at MYC locus. From top to bottom, tracks show the genome scale; Ishikawa contact matrices normalized using the square-root vanilla coverage method for T0, E2 (E), and R5020 (R) treatments; Virtual 4C profiles centered at the MYC TSS, generated from the corresponding Hi-C matrices. Relative interaction is defined as the interaction count at each bin divided by the sum of interaction counts across the entire profiled range; ENCODE CTCF binding sites in Ishikawa cells (ENCSR000BQE; IDR < 0.05); ENCODE RAD21 binding sites in Ishikawa cells (ENCSR000BTU; IDR < 0.05); UCSC gene annotations; ERa and PR ChIP-seq signal (CPM) in Ishikawa cells, with black lines indicating peaks with FDR < 0.05; ERa ChIP-seq signal (CPM) in endometrial tumors from non-users and tamoxifen users, with black lines indicating peaks with FDR < 0.05; H3K27ac ChIP-seq signal (CPM) in non-users and tamoxifen users; and RNA-seq RPKM values in Ishikawa cells treated with control (T0), E2 (E), or R5020 (R).

These findings suggest that overlapping ERa and PR binding regions may contribute to the regulation of NOTCH signaling genes; however, validation through more precise gene expression analyses in tumor samples is required.

#### TADs based approach

In a second approach, we associated the shared ERbs with genes located within the same topologically associated domains (TADs), based on published Hi-C data for Ishikawa cells (27). Ontology analysis of genes located within TADs containing these ERbs revealed a significant enrichment in several pathways involved in inflammation processes, including the Chemokines Receptors Bind Chemokines (R-HSA-380108), Inflammatory Response, TNF-alpha Signaling via NF-kB and IL-6/JAK/STAT3 Signaling pathways (Figure 2b). Genes from the Chemokines Receptors Bind Chemokines and Inflammatory Response pathways were pooled and a GSEA on the endometrial tumor microarray dataset was conducted. This analysis revealed a significant enrichment in non-users compared to tamoxifen users (FDR *≤* 0.1)(Figure 2c), suggesting higher expression of inflammatory genes in the absence of tamoxifen. This subset of genes is distributed across different TADs, with the exception of the families of chemokines (C-X-C subfamily chemokines) and receptors (CCR) that cluster as expected (see supplementary material).

To assess the prognostic value of these pathways in clinical samples, we analyzed RNA-seq data from 394 endometrial adenomas and adenocarcinomas obtained from The Cancer Genome Atlas Program (TCGA) (see Materials and Methods) and found that a higher expression of the Chemokine Receptors Bind Chemokines pathway was significantly associated with improved overall survival (p-value *≤* 0.05; Figure 2d), whereas no significant association was observed for the Inflammatory Response pathway (p > 0.05; Figure 2—Figure supplement 1d).

Ishikawa cells showed a differential expression of several genes within these pathways in response to E2 and tamoxifen treatment, including IL1R1, CDKN1A, SLC11A2, ICOSLG, CXCL11, P2RY2, MYC, ATP2B1 and ITGB8 (Figure 2e). Notably, MYC expression was higher following E2 and tamoxifen treatment than in vehicle-treated cells (Figure 2e) (55).

Among these, the proto-oncogene MYC, a key regulator of inflammation, emerged as a notable candidate deregulated in EC (56–58). The epigenetic landscape of the MYC locus (Figure 2f) showed a cluster of PRbs upstream of the MYC transcription start site (TSS) with high contact frequency. Virtual 4C track shows an increase in interaction in response to hormones. This subset overlaps more with ERbs from non-users than with those from tamoxifen users. There is also an increase in H3K27ac ChIP-seq signal at the MYC locus in tamoxifen users (Figure 2f, Tamoxifen users H3K27ac signal). Reanalysis of RNA-seq data from Ishikawa cells (27) showed significantly increased MYC expression following E2 treatment compared to control (padj *≤* 0.05), with 1 hour tamoxifen treatment also inducing an increase in expression comparable with 1 hour E2 treatment (padj *≤* 0.05) (Figure 2f). Together, these data suggest a complex regulatory landscape involving inflammatory genes and proto-oncogenes such as MYC, with divergent regulatory mechanisms in tamoxifen users and non-users.

### ER occupancy at PR-independent sites in tamoxifen treated tumors

Next, we investigated the genomic proximity of Ishikawa ERbs to the nearest PRbs (Figure 3—Figure supplement 1). Most ERbs were located within 5 kb of a PRbs (Figure 3—Figure supplement 1). To better dissect their potential interactions, we categorized ERbs into two groups: those overlapping PRbs by at least one base pair (ER overlapping PR) and those not overlapping PRbs (*≥* 1 bp from the nearest PRbs) (ER not overlapping PR) (Figure 3a). The former group likely represents loci potentially co-regulated by both ERa and PR, whereas the latter are more likely to be independent of direct PR influence. Our analysis revealed that 732 out of the 1591 Ishikawa ERbs overlapped with PRbs (Figure 3a). To explore whether the regulatory behavior of these two categories differs between tamoxifen users and non-users, we focused on Ishikawa ERbs-both overlapping and not overlapping PRbs-that were also detected in tumors from either of the tamoxifen users or non-users. Motif enrichment analysis on these sites revealed distinct sequence signatures between ERbs overlapping and not overlapping PRbs (Figure 3b). As expected, PGR and AR motifs were only enriched in ERbs overlapping PR. NR1D1, NR1D2 and RORB motifs showed a similar pattern, with a higher enrichment in ERbs overlapping PRbs compared to not overlapping (log2(enrichment ratio) *≈* 1 vs. *≈* 0). In contrast, the PAX motifs PAX1, PAX5 and PAX8 showed the opposite pattern. Motifs of nuclear receptors with response elements similar to the ERE, such as ESR2, ESRRA and PPARG, were also more enriched in ERbs not overlapping PR (log2 enrichment ratio *≈* 2.5–5 vs. *≈* 1–2.5 in ERbs overlapping PR), which is consistent with more frequent direct ERa binding at these regions.

**Figure 3.**
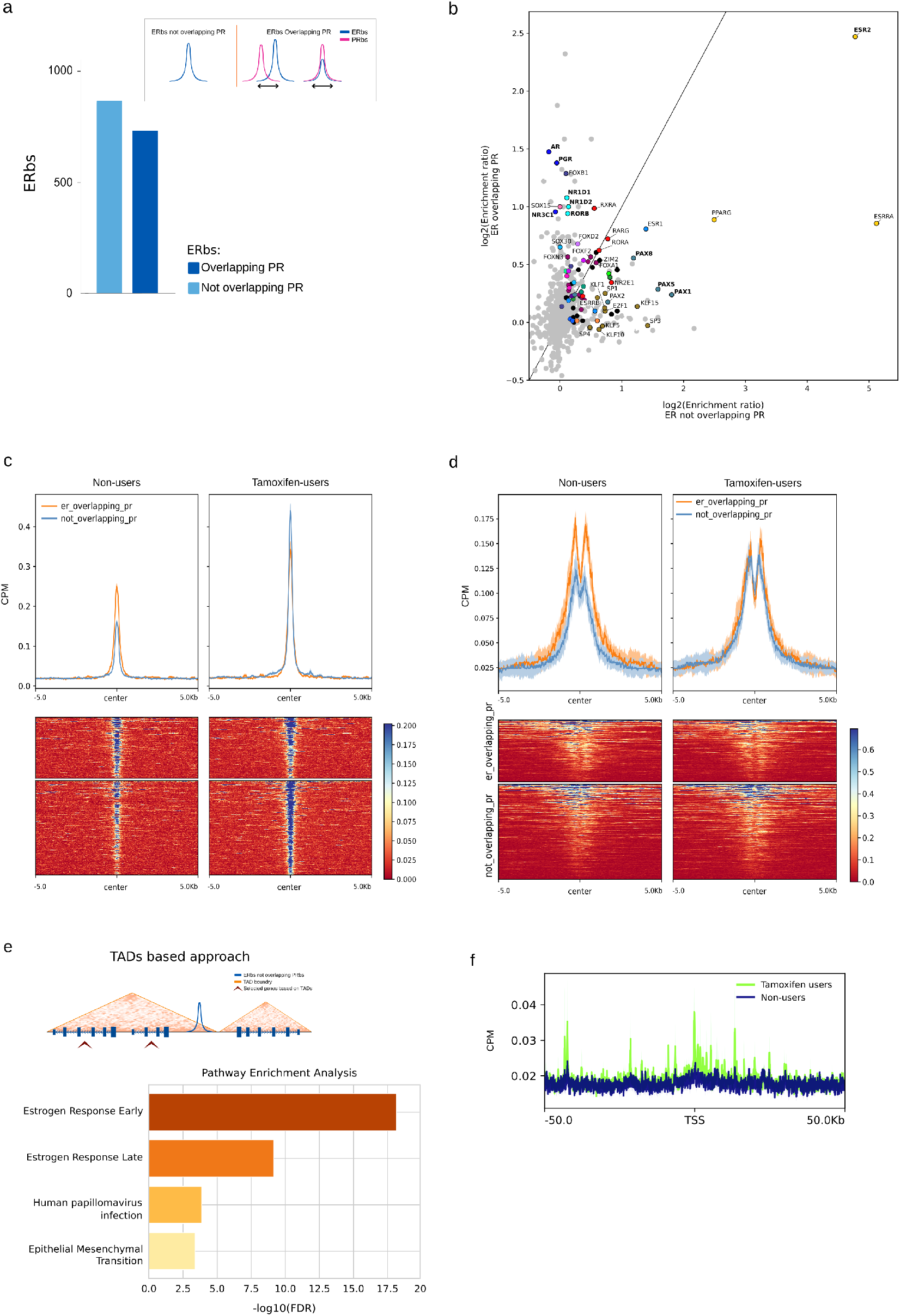
ERbs overlapping PRbs and regulation of Hallmark Estrogen Response Early Pathway. **a.** Number of Ishikawa ERbs that overlap or do not overlap with Ishikawa PRbs. Inset illustrates the criteria used to define both groups. **b**. Scatter plot comparing the log2 enrichment ratio of each motif between ERbs not overlapping PRbs and ERbs overlapping PRbs. Enrichment ratios were calculated with Simple Enrichment Analysis (See Materials and Methods). The diagonal line marks equal enrichment in both groups. Motifs with a significant enrichment (FDR < 0.05) in at least one of the groups are colored, with motifs belonging to the same cluster sharing the same color (Figure 1—Figure supplement 3) and motifs that did not cluster with any other motif are shown in black. Motifs that were not significant in either group are shown in light gray. ZNF motifs were excluded. Labeled motifs were selected manually for visualization, without a predefined selection criterion. Bold labels indicate motifs discussed in the text. **c**. ERa signal distribution in tamoxifen users and non-users. Regions were defined as 10kb windows centered on Ishikawa ERbs (*±*5 kb) that were also identified as ERbs in at least one tumor group (tamoxifen users or non-users), categorized as in (a). Signal intensity represents the median accumulated read count per region, with standard error shown in a lighter tone. **d**. H3K27ac ChIP-seq signal distribution in tamoxifen users and non-users. Regions were defined as in (c). Signal intensity represents the median accumulated read count per region, with standard error shown in a lighter tone. **e**. Representation of the TADs-based approach used to associate binding sites to nearby genes (top). Top pathways enriched in genes located within TADs containing at least one Ishikawa ERbs that does not overlap a PRbs (bottom). **f**. Median of ERa ChIP-seq signal in tamoxifen users and non-users around the transcription start sites (TSS *±*50 kb) of the 200 genes in the Hallmark Estrogen Response Early Pathway. Standa1rd5 error is indicated in a lighter tone.

We next compared ERa ChIP-seq signal between tumors from tamoxifen users and non-users at these two groups of sites. At ERbs not overlapping PRbs, ERa signal was higher in tamoxifen users than in non-users (Figure 3c). At ERbs overlapping PRbs, the pattern was reversed, with a higher ERa signal in non-users (Figure 3c). The H3K27ac signal showed a similar pattern (Figure 3d). In non-users, sites overlapping PR displayed a higher H3K27ac signal than sites not overlapping PR, whereas in tamoxifen users this difference was lost. Together, these results are consistent with a redistribution of ERa from regions overlapping PR toward non-overlapping PR regions in tamoxifen users, accompanied by a reduced enhancer activity at the shared sites.

To identify potential target genes associated with these binding sites, we applied the two approaches described previously. Ontology analysis of genes located within the same TADs as ERbs overlapping PRbs revealed significant enrichment for the Estrogen Response Early pathway (Figure 3e). When we instead considered genes within TADs harboring binding sites shared by both Ishikawa ERbs and PRbs, the Estrogen Response Early pathway remained the most enriched, albeit with lower significance compared to ERbs not shared with PR (Figure 3—Figure supplement 2a). Similar trends were observed with the second approach, which focused on genes proximal to the binding sites: the Estrogen Response Early pathway showed stronger enrichment for ERbs not shared with PRbs (Figure 3—Figure supplement 2b) compared to those shared with PRbs (Figure 3—Figure supplement 2c). Finally, we analyzed the ERa ChIP-seq signal near the transcription start sites (TSSs) of Estrogen Response Early genes in tamoxifen users and non-users (Figure 3f), which showed that the ERa signal around the TSSs of these genes was higher in tamoxifen users compared to non-users.

### Tamoxifen-associated Chromatin Reorganization Influences NRIP1 Expression and Estrogen Response Gene Activity

To determine whether ERa binding near genes within the Estrogen Response Early pathway leads to changes in gene expression, we first performed an ontology analysis of genes differentially upregulated in Ishikawa cells treated with E2 compared to R5020 (padj *≤* 0.05, log2 fold change *≥* 0), using data from our previous study (27). As expected, genes from the Estrogen Response Early pathway were upregulated when treated with estrogen (Figure 4a). We next performed a GSEA analysis of this pathway using microarray expression profiles from tamoxifen users and non-users (23) (Figure 4b). This revealed higher expression of Estrogen Response Early pathway in tamoxifen users compared to non-users (FDR*≤* 0.1). We next evaluated the prognostic value of the expression of the Estrogen Response Early Pathway in the TCGA endometrial cancer cohort. Kaplan-Meier survival analysis of patients with advanced-stage tumors (stage III–IV) showed that a higher expression of the Estrogen Response Early pathway was significantly associated with improved overall survival (p-value *≤* 0.05 Figure 4c).

**Figure 4.**
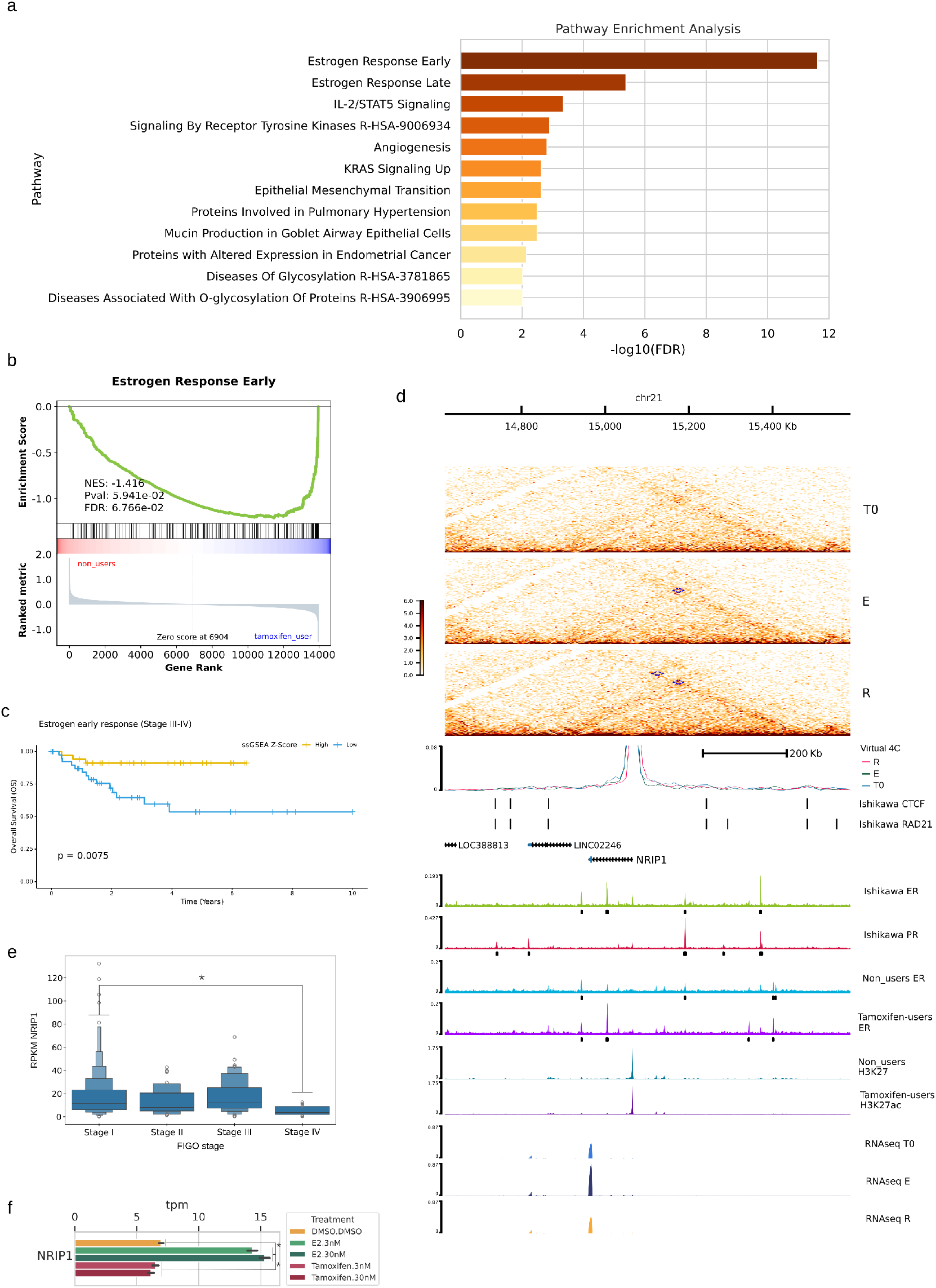
Pathway enrichment and regulatory features associated with NRIP1 and estrogen signaling. **a.** Top pathways enriched among genes upregulated in Ishikawa cells treated with E2 compared with R5020 (padj *≤* 0.05, log2 fold change *≥* 0). **b**. Gene Set Enrichment Analysis (GSEA) of the Hallmark Estrogen Response Early pathway using gene expression profiles from tamoxifen users and non-users. The enrichment profile (green) indicates the correlation with the ranked gene list. The Normalized Enrichment Score (NES) and nominal p-values (nom.p) are shown. **c**. Kaplan-Meier overall survival (OS) curves of FIGO stage III–IV endometrial cancer patients (TCGA data) stratified into high (yellow) and low (blue) pathway expression groups, using the median ssGSEA z-score as a cutoff for the Estrogen Early Response pathway. Log-rank test p-value is reported. **d**. Genomic and epigenetic landscape and gene expression at the NRIP1 locus. From top to bottom, tracks show the genome scale; Ishikawa contact matrices normalized using the square-root vanilla coverage method for R5020 (R) and E2 (E) treatments, with blue dashed lines indicating chromatin loops (FDR < 0.1); Virtual 4C profiles centered at the NRIP1 transcription start site (TSS), generated from the corresponding Hi-C matrices. Relative interaction is defined as the interaction count at each bin divided by the sum of interaction counts across the entire profiled range; ENCODE CTCF binding sites in Ishikawa cells (ENCSR000BQE; IDR < 0.05); ENCODE RAD21 binding sites in Ishikawa cells (ENCSR000BTU; IDR < 0.05); UCSC gene annotations; ERa and PR ChIP-seq signals (CPM) in Ishikawa cells, with black lines indicating peaks with FDR < 0.05; ERa ChIP-seq signals (CPM) from non-users and tamoxifen users, with black lines indicating peaks with FDR < 0.05; H3K27ac ChIP-seq signals from non-users and tamoxifen users; and RNA-seq RPKM values from Ishikawa cells treated with control (T0), E2 (E), or R5020 (R). **e**. NRIP1 expression (RPKM) from RNA-seq data in The Cancer Genome Atlas (TCGA), including 394 endometrial carcinomas. Tumors are grouped by FIGO stage. Asterisks (*) indicate statistically significant differences in expression (padj value *≤* 0.05). **f**. NRIP1 expression (TPM) in Ishikawa cells treated with DMSO, 3nM, and 30nM of estrogen or tamoxifen. Asterisks (*) indicate statistically significant differential expression between treatment groups (padj *≤* 0.05).

Within this pathway, the Nuclear Receptor Interacting Protein 1 (NRIP1) gene emerged as an interesting candidate for further investigation (Figure 4). NRIP1 encodes a nuclear co-regulator that modulates the transcriptional activity of various transcription factors, including ER. Its differential epigenetic landscape between tamoxifen users and non-users suggests altered regulation of endometrial tumors (Figure 4d). Hi-C chromatin conformation analysis revealed a differential loop (highlighted by blue dashed lines) present in R5020-treated, but not in E2-treated, Ishikawa cells. This loop connects two CTCF binding sites. In addition, a virtual 4C analysis centered on the NRIP1 TSS showed no changes in the contact profile.

Expression profiling of NRIP1 in Ishikawa cells showed a downregulation under R5020 treatment compared to E2, and an upregulation when comparing E2 to untreated conditions (padj *≤* 0.05) (Figure 4d, 3 last tracks). Similar epigenetic features were also observed for the genes: Growth Regulated by Estrogen in Breast Cancer 1 (GREB1), Cyclin A1 (CCNA1) and Dehydrogenase/reductase 3 (DHRS3) from the Estrogen Response Early pathway (Figure 4—Figure supplement a,b,c). To assess NRIP1 expression in clinical samples, we analyzed the TCGA endometrial cancer cohort (Figure 4e) and found that it was significantly lower in stage IV than in stage I tumors (padj *≤* 0.05). In the same cohort, NRIP1 and ESR1 expression were positively correlated (partial correlation = 0.47, adjusted for age and FIGO stage; Figure 4—Figure supplement 1d), in line with previous reports that NRIP1 expression correlates with ERa levels (59). The lower NRIP1 expression in advanced tumors may therefore reflect the reduced ERa expression typically observed at later stages of endometrial cancer. Consistent with NRIP1 being estrogen responsive, its expression in Ishikawa cells was significantly higher after E2 treatment than after tamoxifen treatment or in controls (padj *≤* 0.05; Figure 4f).

Collectively, these results suggest that the Estrogen Response Early Pathway plays a pivotal role in shaping endometrial response to tamoxifen. Prolonged tamoxifen treatment may enhance ERa binding at sites with limited PR co-regulation, leading to upregulation of the Estrogen Response Early pathway, influencing key genes such as NRIP1, and contributing to a distinct genomic and transcriptional landscape in tamoxifen-associated endometrial cancer compared to tumors from non-users.

## Discussion

Our analysis revealed that the ERa binding profile in tamoxifen users closely mirrors the ERa landscape observed in estrogen-treated Ishikawa cells, whereas the ERa profile of non-users aligns more closely with the PR binding pattern in the same cell line. This shift is consistent with previous observations by (22, 23), which demonstrated a distinct ERa binding profile in tamoxifen-associated endometrial tumors. Importantly, PGR expression did not differ substantially between tamoxifen-associated and non-associated tumors (Fig. 5—Figure supplement 1). This suggests that the observed differences in the relationship between ERa and PR binding are unlikely to be explained simply by altered PGR transcription. However, differences in PR protein abundance, post-translational regulation, or receptor activity cannot be excluded. Prior studies have established that PR can modulate ERa activity and down-stream gene expression in breast cancer (24) and that the coordinated binding of ERa, PR, and PAX2 regulates transcriptional programs implicated in endometrial carcinogenesis (27). In this context, our findings propose that tamoxifen disrupts PR-mediated regulation of ERa, resulting in alternative transcriptional programs that may contribute to cancer progression.

**Figure 5.**
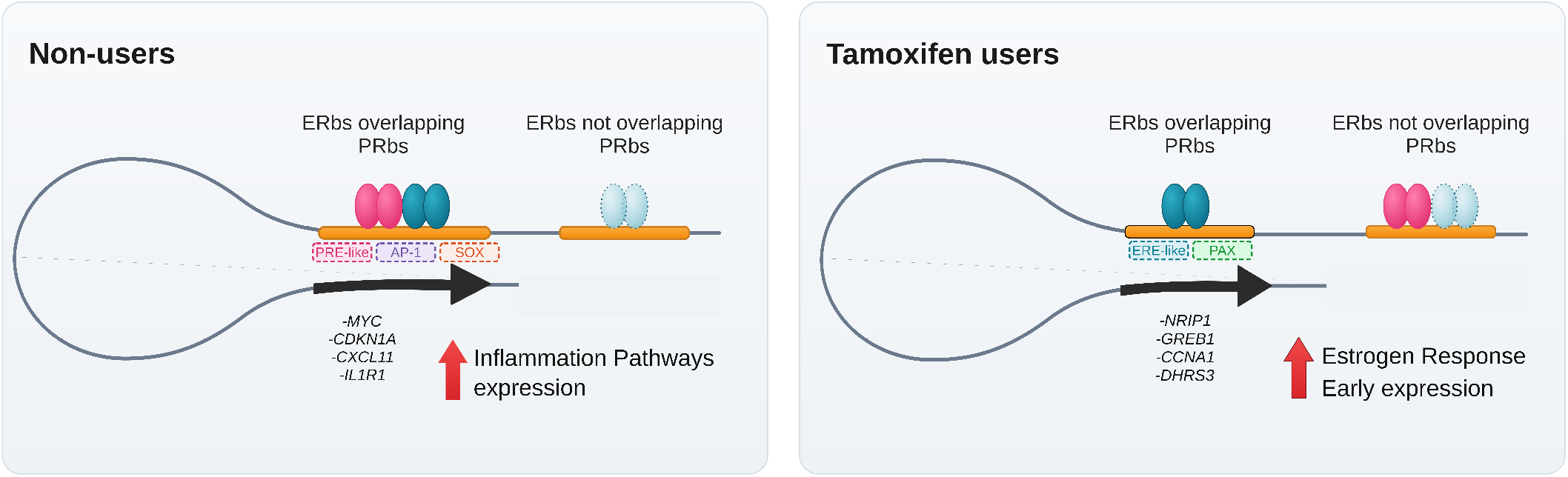
Proposed model of tamoxifen-associated changes in ERa-PR regulation in endometrial cancer. Schematic of ERa (blue) and PR (pink) binding at regulatory regions in non-users (left) and tamoxifen users (right). Darker ERa indicates stronger ChIP-seq signal. Dashed boxes show motifs enriched at each group of ERbs. In non-users, ERbs overlapping PRbs are associated with inflammatory pathway expression (red arrow). In tamoxifen users, ERa shifts to non-overlapping sites associated with Estrogen Response Early expression. Listed genes are examples of each pathway.

Motif enrichment analysis of non-users ERbs shared with PRbs revealed enrichment of PGR motifs, together with NR3C1 and AR motifs, whose response elements share sequence similarities with those recognized by PR. In contrast, ESR1 and ESR2 motifs were not enriched, whereas motifs from the AP-1 family were significantly enriched. This pattern raises the possibility that ERa occupancy at these sites involves interactions with other transcription factors, including AP-1, which have been shown to mediate estrogen responsiveness (60). However, motif enrichment alone cannot establish the mechanism of ERa recruitment or demonstrate PR occupancy in tumors. The enrichment of SOX family motifs is also consistent with previous reports of SOX motif enrichment at ERa binding sites in endometrial cancer (52). Further characterization of the differential binding patterns revealed that ERa binding sites uniquely shared with Ishikawa PR sites in non-users were enriched in pathways related to inflammatory responses, which correlated with increased expression of the associated genes. Among inflammatory genes, overexpression of c-Myc has been reported in over 70% of endometrial tumors (57) and its expression correlates with disease prognosis (58). Prior studies indicate that PR can act as a negative regulator of MYC transcription in endometrial cancer (56). In view of this, the presence of PRbs that overlap non-user ERa sites at the MYC locus is consistent with the possibility of PR-associated regulation of this gene in tumors from non-users. In Ishikawa cells, MYC expression was significantly upregulated following both estradiol (E2) and tamoxifen treatment, although it is important to note that short-term in vitro exposure to tamoxifen does not fully replicate the long term treatment in patients.

The anti-inflammatory effects of progesterone, mediated through PR, are well documented in the endometrium, including in endometrial adenocarcinomas (61). Our results are consistent with a model in which PR, acting in concert with ER, regulates inflammatory pathways in non-users, and loss of this regulatory axis may contribute to tumor development (Figure 5). By contrast, tamoxifen-associated EC may engage alternative pathways not typically modulated by PR, such as PI3K signaling (62). These observations underscore the need for further investigation to delineate how tamoxifen-induced shifts in ERa cistromes reprogram transcriptional networks, particularly those governing inflammatory signaling, and how these changes influence endometrial tumor biology.

In tamoxifen users, ERa binding shifts toward sites that do not overlap Ishikawa PRbs. Motif enrichment analysis revealed distinct sequence signatures between ERa sites overlapping and not overlapping PRbs. PGR, AR, NR1D1, NR1D2 and RORB motifs were more enriched at overlapping sites, whereas PAX family motifs and motifs of nuclear receptors with response elements similar to the estrogen response element, including ESR2, ESRRA and PPARG, were more enriched at non-overlapping sites. These differences suggest that the two groups of ERbs have distinct potential transcription factor recruitment profiles, although motif enrichment alone cannot establish direct binding. Together with the differences in ERa and H3K27ac signal between the two groups, these findings support the possibility that tamoxifen-associated changes in ERa occupancy involve a redistribution between regulatory regions with distinct sequence and chromatin features. To investigate the potential functional implications of this redistribution, we analyzed genes located within the same TADs as ERbs not overlapping PRbs. This analysis revealed enrichment of the Estrogen Response Early pathway, which was also more highly expressed in tamoxifen-associated tumors. The ERa co-regulator NRIP1 is a key component of this pathway, and its expression is known to inhibit ERa activity in mammary cells (63). In endometrial cancer, NRIP1 has been identified as a frequently mutated gene in microsatellite-unstable tumors, and its silencing was associated with altered expression of genes within the estrogen-receptor network (64). Recently, epigenomic analysis on the human endometrial stromal cell line tHESC identified NRIP1 as one of the genes upregulated by E2, presenting three ERbs interacting with its promoter (approximately 60kb, 200kb and 270kb away from the TSS) (65). Only one (+60kb) of these ERbs was identified in the epithelial adenocarcinoma cell line Ishikawa treated with E2. Non-users endometrial tumors have not presented these ERbs at all, while tamoxifen users share all of them. This observation could reflect the presence of stromal cells in the tamoxifen user tumor samples, although other causes cannot be excluded.

A major limitation is the lack of in vivo PR cistrome data in endometrial tumors, and particularly in tamoxifen-associated cases. The Ishikawa cell line was selected as a reference because, to our knowledge, they are the only endometrial model with matched ERa/PR cistrome data, but it may not fully capture the complexity of primary tumors. Ishikawa cells cannot model the heterogeneity of tumor subtypes or the stromal–epithelial interactions that shape receptor activity in vivo. In addition, the tumor ChIP-seq data come from a small number of samples (6 non-users and 9 users), and gene expression results from the microarray data showed no individual gene was significantly differentially expressed. Our findings are therefore correlative and exploratory, and require validation with functional experiments in primary tumor material. Nevertheless, the convergence of our findings with prior mechanistic studies supports the hypothesis that tamoxifen reshapes the ER–PR regulatory axis, highlighting the need for future studies using primary tumor material to define PR-dependent modulation of ERa under tamoxifen exposure.

These findings support a model in which prolonged tamoxifen exposure reshapes the ERa–PR regulatory axis, leading to reduced PR-mediated control of inflammatory pathways and enhanced ER-driven activation of the Estrogen Response Early program (Figure 5). These alternative transcriptional states may influence tumor behavior and progression. This exploratory analysis provides a starting point for studying how long-term SERM treatment may alter ERa–PR regulation in the endometrium, with potential implications for prognosis and targeted therapy development.

## Data Availability

ERa and H3K27ac ChIP-seq data and microarray data from endometrial cancers from tamoxifen users and non-users, as reported by (22, 23) are accessible in Gene Expression Omnibus (GEO) under accession number GSE94524. ERa and PR ChIP-seq data along with RNA-seq data from Ishikawa cells treated with E2 and R5020 are available under accession number GSE139398. RNA-seq data from Ishikawa cells treated with tamoxifen and E2 (55) can be accessed via GEO accession number GSE115608. CTCF and RAD21 binding sites in Ishikawa cells, used in omics track figures, are available in ENCODE under accession number ENCSR000BQE and ENCSR000BTU, respectively. RNA-seq data for human endometrial cancer samples were obtained from The Cancer Genome Atlas (TCGA) under project TCGA. Processed data and custom scripts for virtual 4C, TCGA samples analysis and RNA-seq differential expression analysis are accessible at Figshare under reference number 28840463 (https://doi.org/10.6084/m9.figshare.28840463). For information about previously published datasets used see Table 1.

**Table 1.**
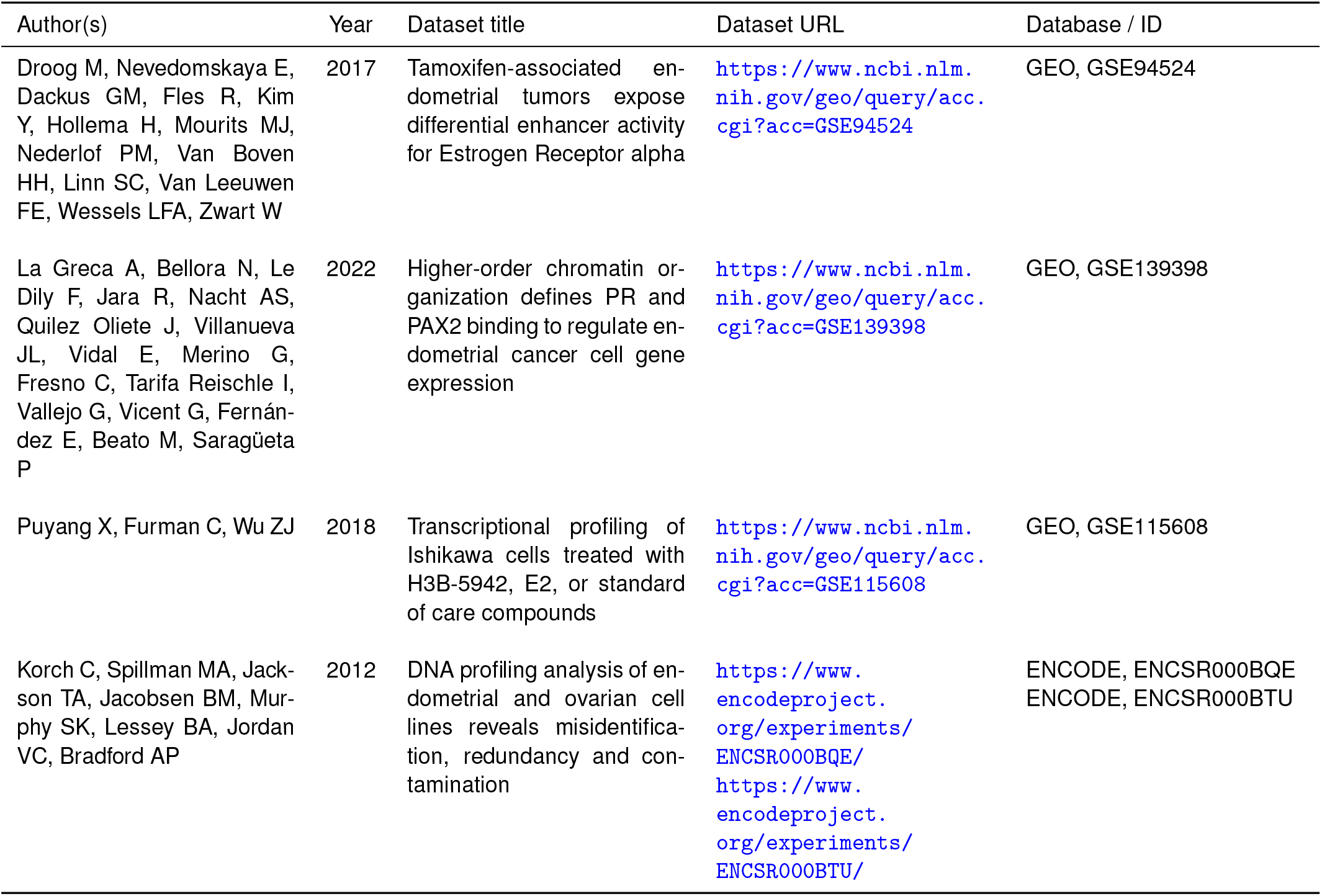
Previously published datasets used in this study.

## Author contributions

**Luciana Ant:** Conceptualization, Data curation, Formal analysis, Investigation, Methodology, Visualization, Writing – original draft, Writing – review and editing; **Nicolás Bellora:** Data curation, Formal analysis, Methodology, Supervision; **Carina Fariñas:** Formal analysis, Investigation, Methodology, Visualization, Writing – original draft; **Alejandro La Greca:** Formal analysis, Writing – review and editing; **François Le Dily:** Formal analysis; **Patricia Saragüeta:** Conceptualization, Formal analysis, Funding acquisition, Investigation, Project administration, Supervision, Writing – original draft, Writing – review and editing.

## Ethics Statement

The present study relies exclusively on publicly available, open-source data. As no new experiments were conducted on human subjects or animals, and no identifiable data was used, this work did not require ethical approval or review by any committee.

## ACKNOWLEDGEMENTS

Generative AI was used to enhance the clarity, grammar, and style of the text. All scientific content, data interpretation, and conclusions remain the sole responsibility of the authors. We are grateful to members of the Saragüeta laboratory for help and suggestions, to Gabriela Diessler for the support and Irene Singer for the graphic design and illustration of Figure 1a.

## Figures

**Figure 1—Figure supplement 1.**
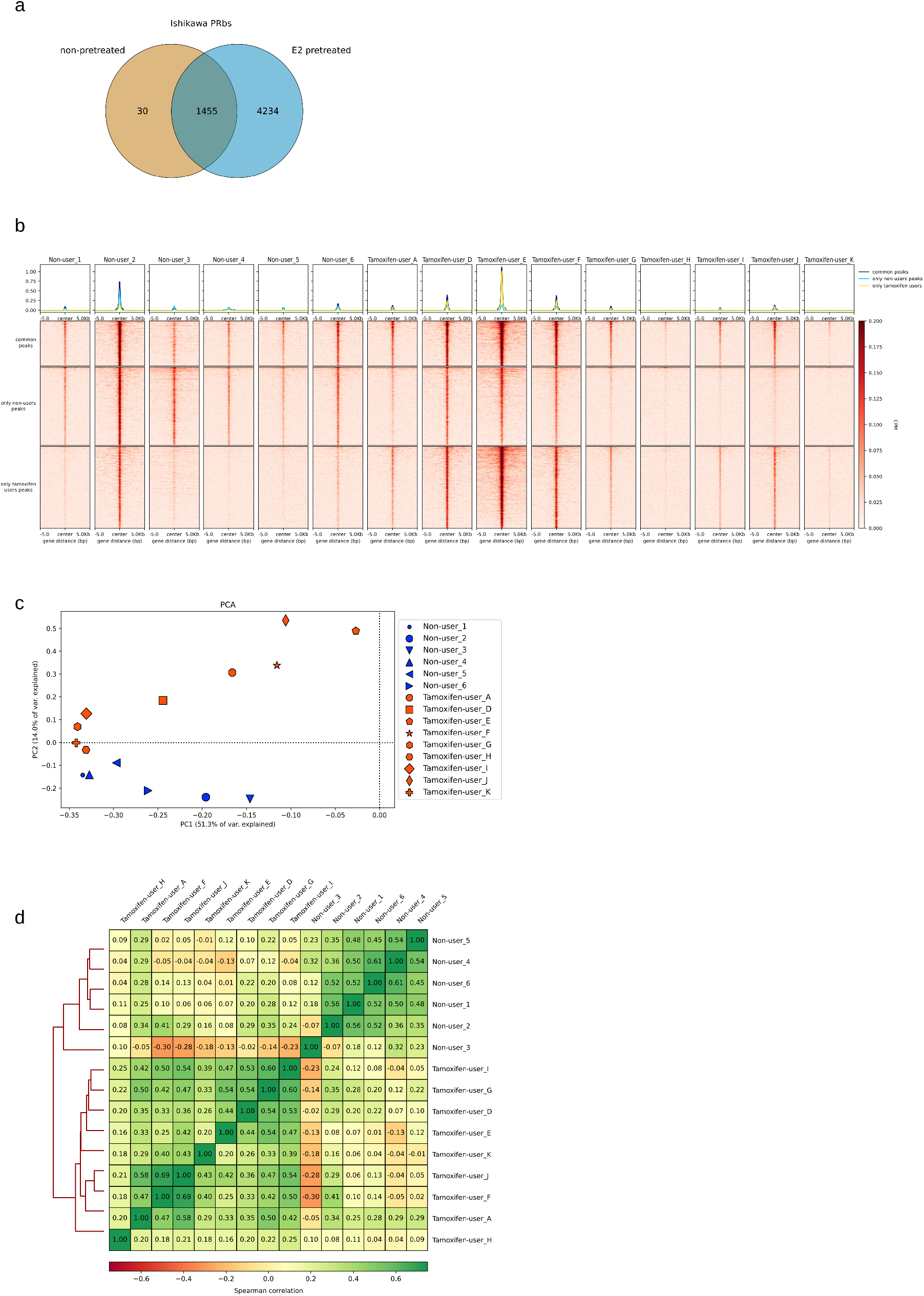
**a**. Venn diagram showing the overlap of PR binding sites (PRbs) in Ishikawa cells pretreated with E2 for 12 hours and those without pretreatment. The intersection represents PRbs shared between both conditions. **b**. Distribution of ERa ChIP-seq signal in tumor samples from tamoxifen users and non-users. Regions were defined as 10 kb windows centered around ERa binding sites (*±*5 kb) identified in tamoxifen users, non-users, or in both groups (common sites). Top: median ERa ChIP-seq signal (CPM) across each ERbs group in tamoxifen users and non-users samples. Bottom: heatmap showing ERa ChIP-seq signal intensity across all samples. **c**. Principal component analysis (PCA) based on ERa ChIP-seq signal across all tamoxifen-user and non-user samples. PC1 is plotted against PC2. **d**. Spearman correlation and hierarchical clustering of tamoxifen-user and non-user tumor samples based on ERa ChIP-seq signal intensity around ERa binding sites.

**Figure 1—Figure supplement 2.**
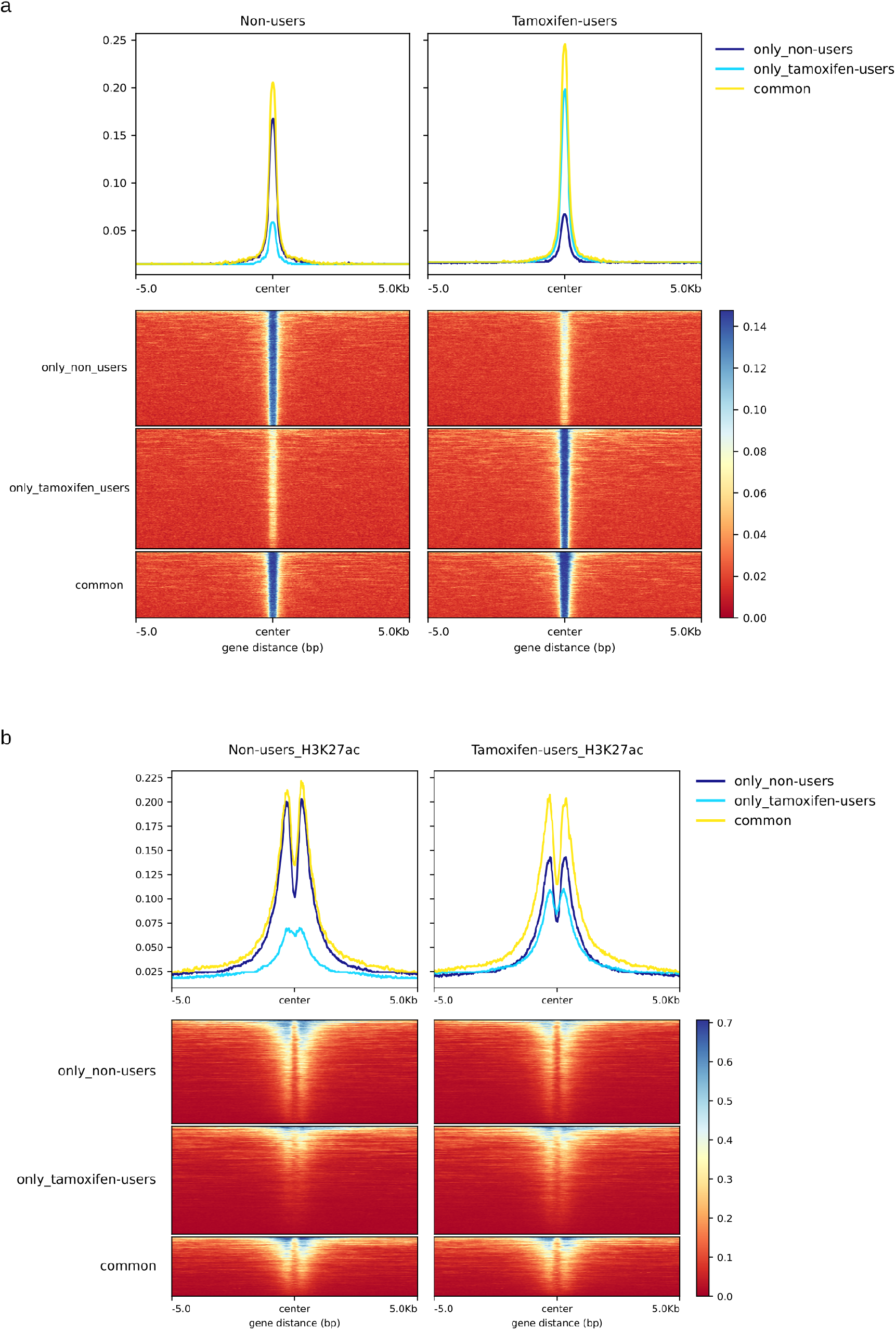
ERa and H3K27ac signal distribution in tumors from tamoxifen users and non-users. **a**. Regions were defined as 10kb windows centered on ERa binding sites (*±*5 kb) identified in tamoxifen users, non-users, or in both groups (common sites). Top panel: median accumulated ERa ChIP-seq signal (CPM) across each ERbs group in tamoxifen users and non-users. Bottom panel: Heatmap showing ERa ChIP-seq signal intensity across individual regions. **b**. Same analysis as in (a), showing H3K27ac ChIP-seq signal in tumors from tamoxifen users and non-users. Regions were defined as in (a).

**Figure 1—Figure supplement 3.**
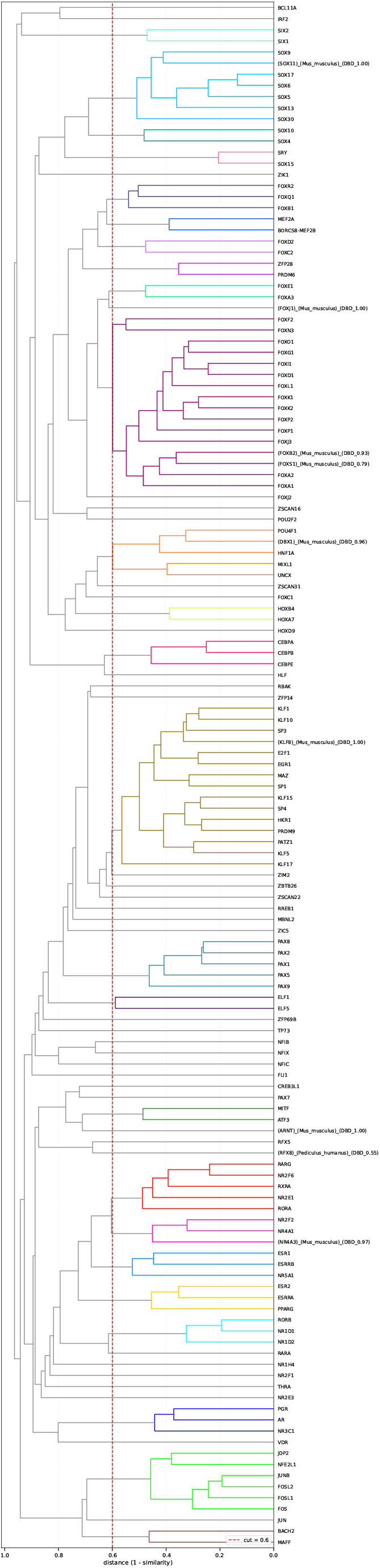
Hierarchical clustering of enriched transcription factor motifs. Dendrogram of the 139 motifs (without including ZNF motifs) that reached FDR *≤* 0.05 in at least one of the three groups (ERa sites from tamoxifen non-users overlapping Ishikawa PR sites, Ishikawa ERa sites not overlapping PR, and Ishikawa ERa sites overlapping PR). Motifs were clustered by the genomic overlap of their predicted sites (see Materials and Methods). The dashed line marks the distance cutoff of 0.6 used to define motif clusters. 37 motifs did not cluster with any other motif at this cutoff and remained as individual (unclustered) motifs. Motif names in parentheses indicate CIS-BP entries inferred from a non-human or paralogous TF.

**Figure 2—Figure supplement 1.**
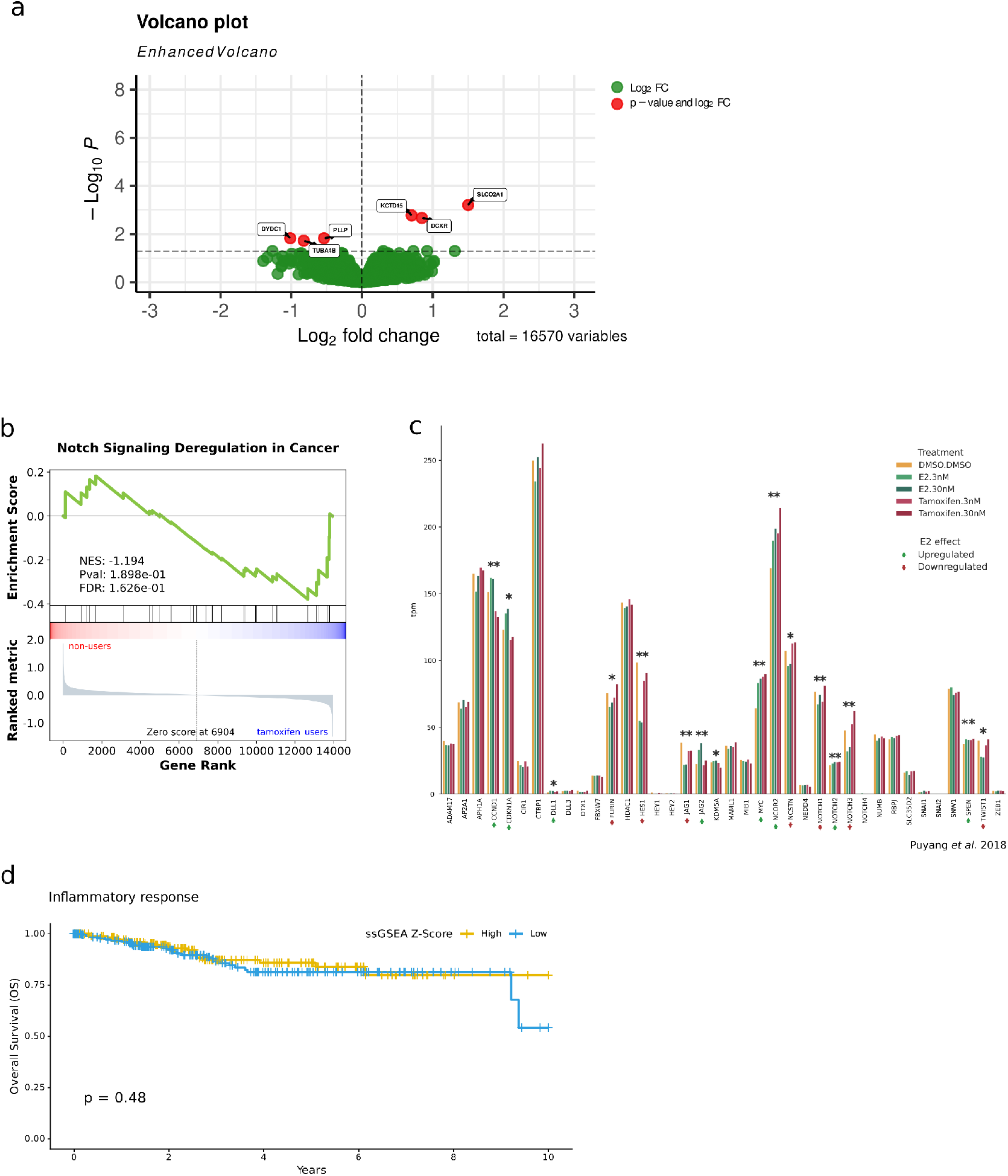
Notch signaling pathway analysis in tamoxifen-treated samples and Ishikawa cells. **a**. Volcano plot depicting the results of differential expression analysis comparing gene expression profiles between tamoxifen users and non-users (N=16570 genes). The dashed horizontal line indicates a padj threshold of 0.05. Genes highlighted in red show a padj < 0.05. **b**. Gene set enrichment analysis (GSEA) of the Notch Signaling Deregulation in Cancer pathway using gene expression profiles from tamoxifen users and non-users. The enrichment profile (green) illustrates the correlation of gene expression with the ranked gene list. Normalized enrichment scores (NES) and nominal p-values (nom. p) are shown. **c**. Expression levels (TPM) of genes from the Notch Receptor Signaling and Notch Signaling Deregulation in Cancer pathways in Ishikawa cells treated with control (DMSO), tamoxifen (3nM and 30nM), or estrogen (3nM and 30nM) (E2.3nM and E2.30nM). One asterisk (*) indicates significant differential expression between E2 (30nM) and DMSO (padj *≤* 0.05). Two asterisks (**) indicate significant differential expression between E2 (30nM) and DMSO and tamoxifen (30nM) and DMSO (Padj). KDM5A is the only gene that presents a differential expression between tamoxifen (30nM) and DMSO but not between E2 (30nM) and DMSO (padj *≤* 0.05). **d**. Kaplan-Meier overall survival (OS) curves of endometrial cancer patients (TCGA data) stratified into high (yellow) and low (blue) pathway expression groups, using the median ssGSEA z-score as a cutoff for the Inflammatory Response pathway. Log-rank test p-value is reported.

**Figure 3—Figure supplement 1.**
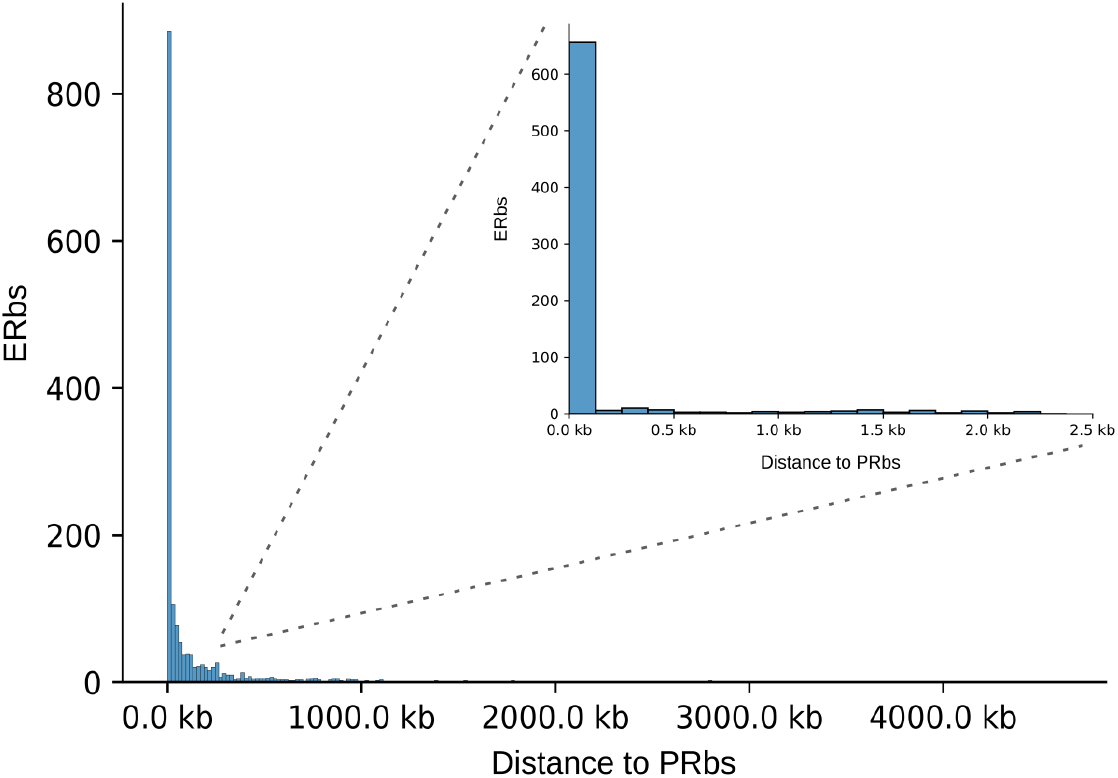
Distance between ERa and PR binding sites in Ishikawa cells. Histogram of the distribution of Ishikawa ERbs according to their distance from the nearest Ishikawa PRbs. The inset highlights the subset of ERbs located within 2.5 kb of a PRbs.

**Figure 3—Figure supplement 2.**
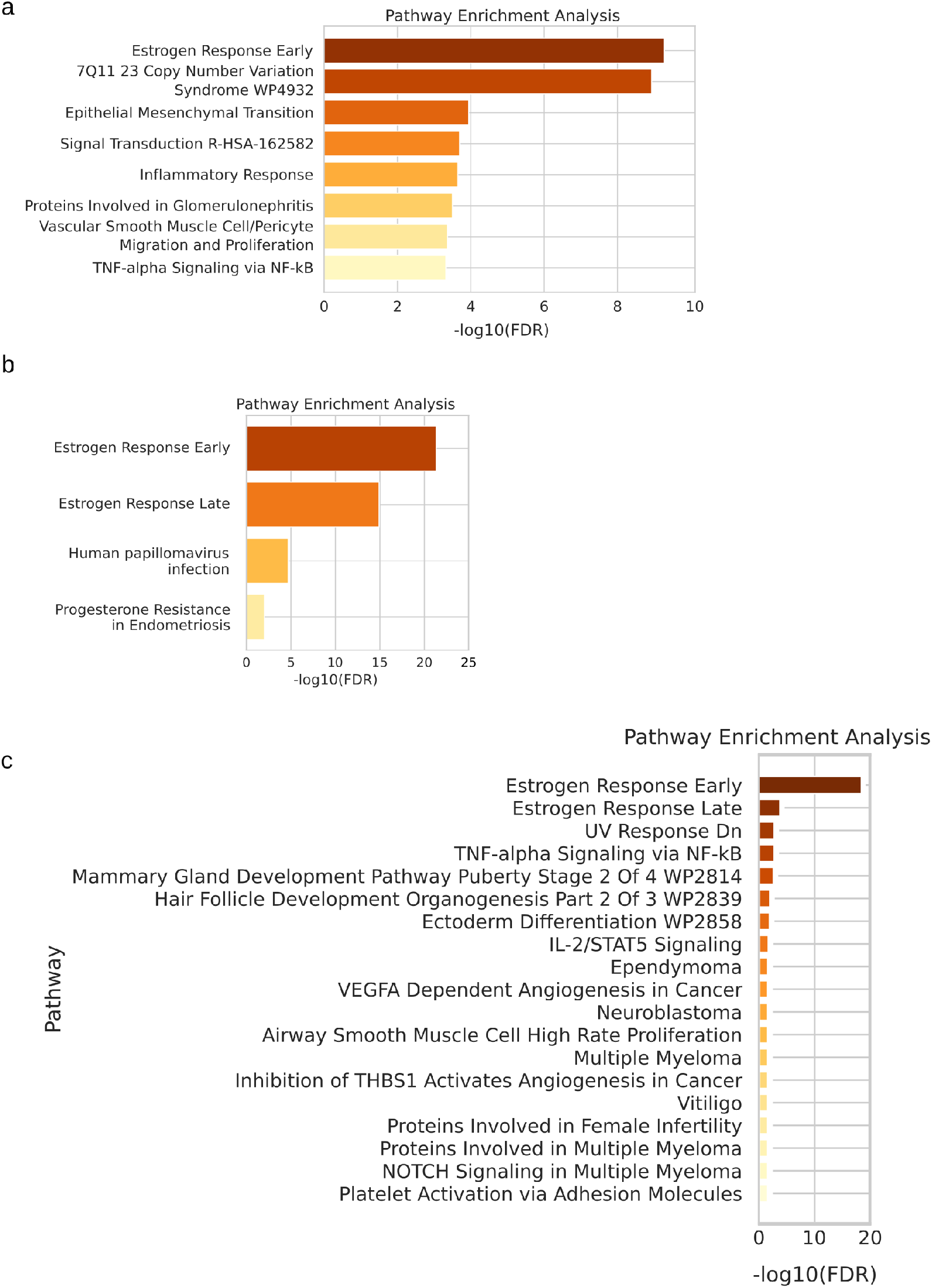
Pathway enrichment associated with ERa and PR binding sites co-occupancy in Ishikawa cells and tumor samples. **a**. Top enriched pathways for genes located in TADs containing at least one Ishikawa ERbs that overlaps with a PRbs and is also present as an ERbs in either tamoxifen users or non-users. **b**. Top enriched pathways for genes located near Ishikawa ERbs that do not overlap with PRbs but are also present as ERbs in either tamoxifen users or non-users. **c**. Top enriched pathways for genes located near Ishikawa ERbs that overlap with PRbs and are also present as ERbs in either tamoxifen users or non-users.

**Figure 4—Figure supplement 1.**
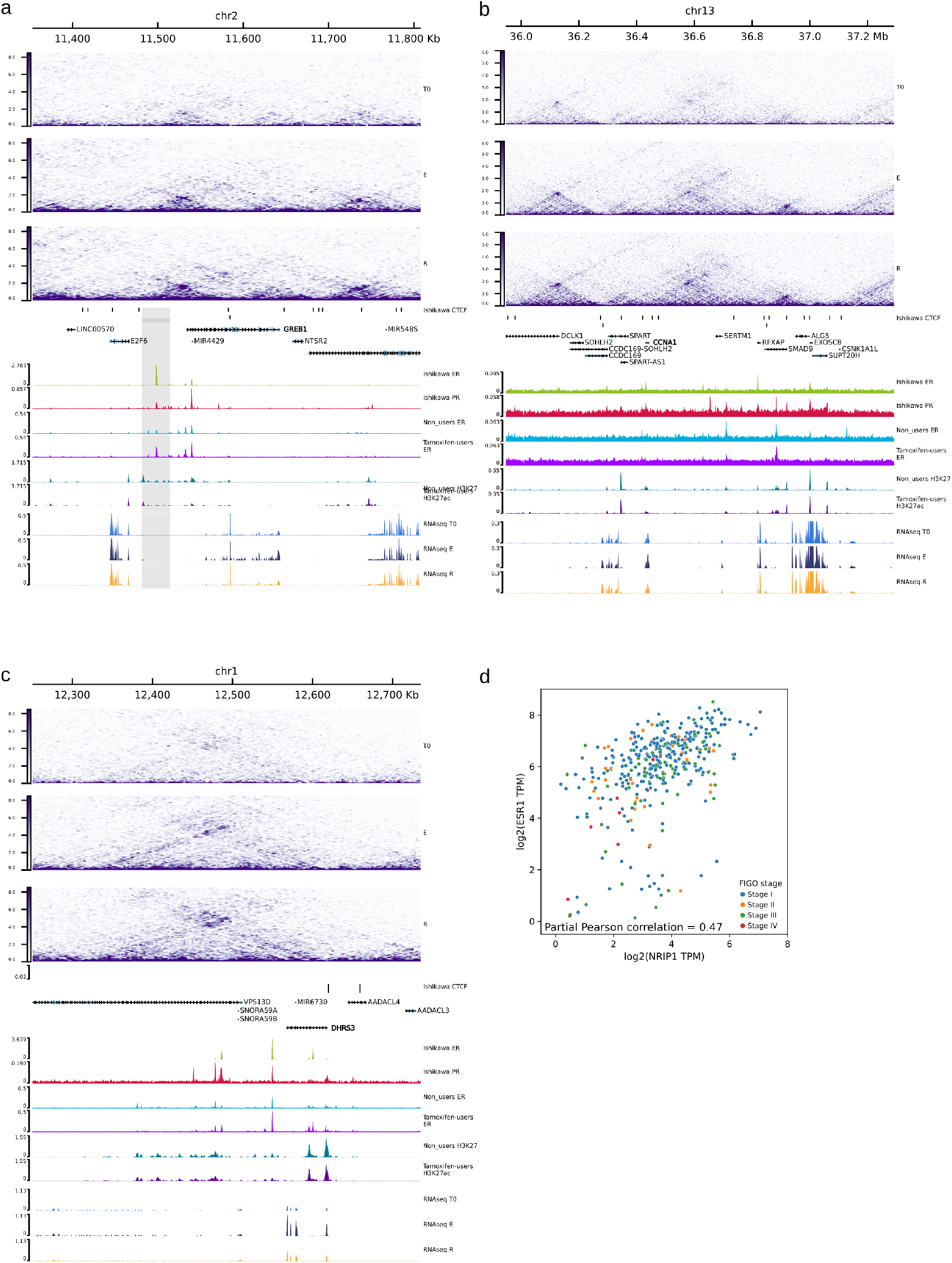
Epigenetic landscape and gene expression at the GREB1 (a), CCNA1 (b), and DHRS3 (c) loci. From top to bottom, tracks show: genome scale; contact matrices from Ishikawa cells treated with control (T0), E2 (E), or R5020 (R), normalized using the square-root vanilla coverage method; ENCODE CTCF binding sites in Ishikawa cells (ENCSR000BQE; IDR < 0.05); UCSC gene annotations; ERa and PR ChIP-seq signals (CPM) from Ishikawa cells; ERa ChIP-seq signals (CPM) from tamoxifen users and non-users; H3K27ac ChIP-seq signals (CPM) from tamoxifen users and non-users; and RNA-seq RPKM values from Ishikawa cells treated with control (T0), E2 (E), or R5020 (R). **d**. ESR1 and NRIP1 expression in the TCGA cohort. Gene expression values (TPM) were log2-transformed. Each point represents a TCGA sample, colored according to FIGO stage. Partial correlation between ESR1 and NRIP1 expression, controlling for age and FIGO stage, is shown. The 95% confidence interval for the correlation coefficient is [0.39, 0.54], with an associated p-value of 1.5 *×* 10^*−*22^.

**Figure 5—Figure supplement 1.**
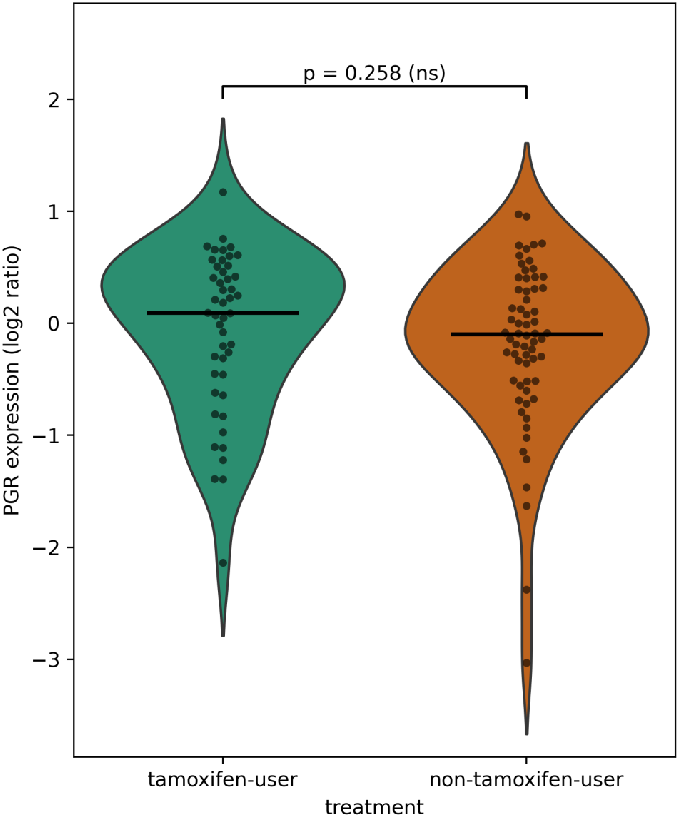
Progesterone receptor expression in endometrial tumors from tamoxifen users versus non-users, measured by microarray as log2(tumor/reference pool). Each dot represents individual samples for each group. The p-value for a two-sided Mann-Whitney U test is shown.

